## Supplementary Figure S1 for "The stage specific plasticity of descending modulatory controls in a rodent model of cancer induced bone pain"

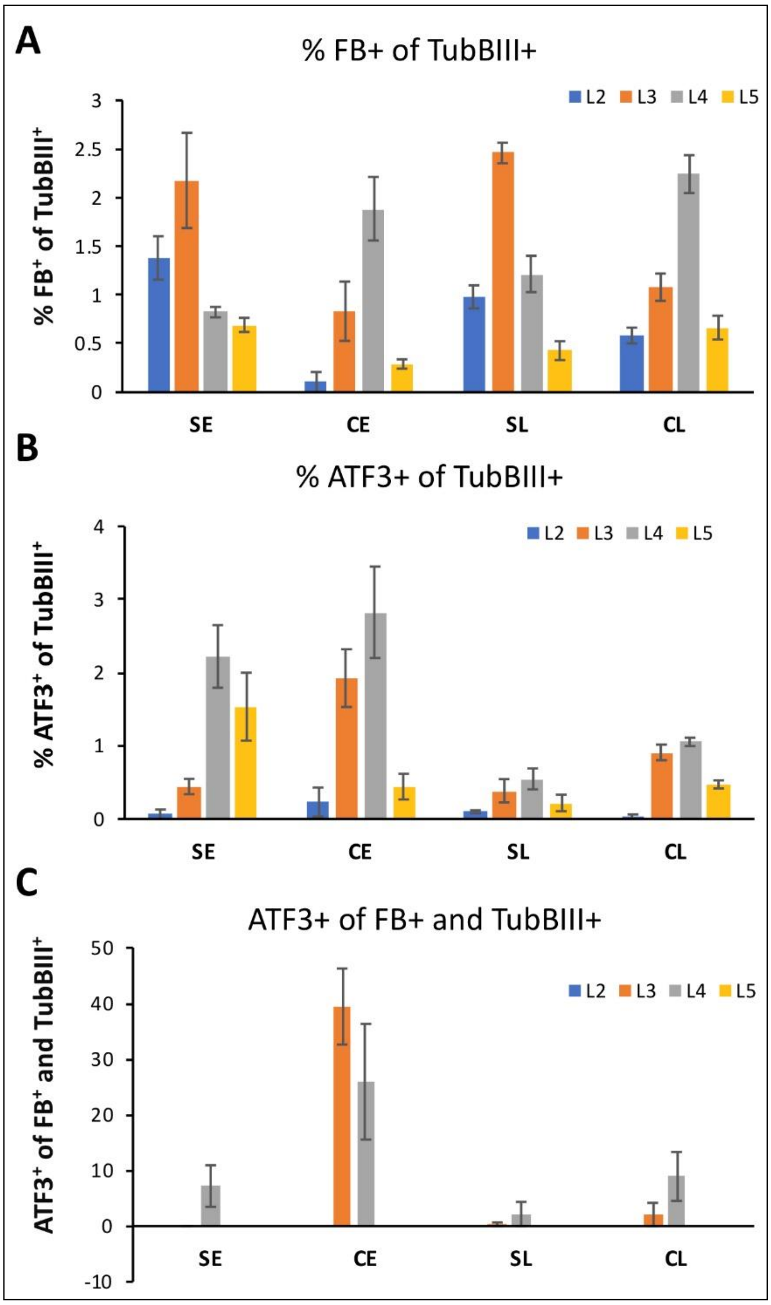


**Figure S1: Detailed Atf3 quantification in separate dorsal root ganglia. (A)** Total number of Fast Blue (FB) traced tibial afferents within ipsilateral L2-5 DRG analysed as a percentage of all neurons (Tubullin-βIII) therein. On average 100.6 ± 15.7 L2-5 DRG neurons innervate tibia. No FB positivity was noticed in the contralateral lumbar DRG (not shown). **(B)** Quantification of all Atf3$^+$ afferents within ipsilateral L2-5 DRG, analysed as a percentage of all neurons (Tubullin-βIII) therein in cancer early (CE, day 7/8) and cancer late (CL, day 14/15) stage groups with corresponding sham groups (early – SE and late – SL). **(C)** Quantification of Atf3+ afferents within ipsilateral L2-5 DRG analysed as a percentage of all FB traced neurons. On average 4-20 10 µm sections were counted per DRG. Data represent the mean ± SEM (n = 3, 'n' denotes a separate animal).
